# Avoidance of toxic misfolding and protein stability do not explain the sequence constraints of highly expressed proteins

**DOI:** 10.1101/168963

**Authors:** Germán Plata, Dennis Vitkup

## Abstract

The avoidance of cytotoxic effects associated with protein misfolding has been proposed as a dominant constraint on the sequence evolution and molecular clock of highly expressed proteins. Recently, Leuenberger *et al.* developed an elegant experimental approach to measure protein thermal stability at the proteome scale. The collected data allow us to rigorously test the predictions of the misfolding avoidance hypothesis that highly expressed proteins have evolved to be more stable, and that maintaining thermodynamic stability significantly constrains their evolution. Notably, careful re-analysis of the Leuenberger *et al.* data across four different organisms reveals no substantial correlation between protein stability and protein abundance. Therefore, the key predictions of the misfolding toxicity and related hypotheses are not supported by available empirical data. The data also suggest that, regardless of protein expression, protein stability does not substantially affect the protein molecular clock across organisms.

---

A fundamental and long-standing question in molecular evolution is what determines protein sequence constraints, or the rate of the protein molecular clock (Zuckerkandl and Pauling 1965; Zhang and Yang 2015). Proteins from the same species accumulate substitutions at rates that span several orders of magnitude, and the causes of such variability have been widely debated (Koonin and Wolf 2010). Analyses of high-throughput genome-scale data consistently showed that protein evolutionary rates are strongly anticorrelated with their corresponding expression and abundance levels (Pal et al. 2001; Pal et al. 2006). This relationship, often referred to as the E-R (<u>E</u>xpression-evolutionary <u>R</u>ate) anticorrelation (Zhang and Yang 2015), explains up to a third of the variance in molecular clock rates across proteins (Pal et al. 2006; Drummond and Wilke 2008). Among possible explanations of the E-R anticorrelation is the popular hypothesis that highly expressed proteins evolve slowly to avoid mistranslation-induced (Drummond and Wilke 2008) or spontaneous (Yang et al. 2010) protein misfolding. According to this hypothesis, misfolded proteins are toxic to cells and therefore reduce fitness. As highly abundant proteins have the potential to produce more misfolded proteins compared to proteins with low abundance, their sequences should be under stronger evolutionary constraints to increase protein stability (Drummond and Wilke 2008; Zhang and Yang 2015). Thus, a key prediction of the misfolding avoidance hypothesis is that highly expressed proteins should be more thermodynamically stable than proteins expressed at low levels, and that selection against protein misfolding should significantly constrain their sequence evolution (Cherry 2010; Serohijos et al. 2012; Serohijos et al. 2013).

Previously (Plata et al. 2010), based on a small set of proteins available in the proTherm database (Bava et al. 2004), we did not detect any significant correlation between protein expression and thermodynamic stability. Furthermore, to empirically test the misfolding hypothesis, we expressed wild type (WT) and destabilized mutant versions of the LacZ protein in *Escherichia coli.* This analysis demonstrated that the corresponding fitness effects were primarily related to the cost of gratuitous protein production and not to misfolding toxicity (Plata et al. 2010). Similar experiments in yeast by Kafri *et al.* (2016) using WT and destabilized versions of GFP, also showed that misfolded protein toxicity plays a relatively minor role in explaining the fitness cost behind the E-R anticorrelation.

As the aforementioned results have been obtained using small sets of proteins, additional tests involving large datasets across diverse organisms are essential. Recently, Leuenberger *et al.* (2017) measured the thermal stability of thousands of proteins across two bacteria (E. *coll* and *Thermus thermophllus)* and two eukaryotes *(Saccharomyces cerevisiae* and *Homo sapiens).* The unprecedented size of this dataset, measured directly in the cellular matrix, provides a unique opportunity to empirically test the misfolding toxicity hypothesis. Using protein melting temperatures (T_m_) from *E. coli*, Leuenberger *et al.* concluded that highly abundant proteins are stable because they are evolutionarily designed to tolerate translational errors (Leuenberger et al. 2017), supporting the misfolding toxicity avoidance hypothesis. The authors reached their conclusion based on different abundances of *E. coli* proteins separated into three bins according to their thermal stability (Figure 3I in Leuenberger *et al*.), but did not perform similar analyses for the remaining three species. Notably, analyses of arbitrarily binned data may often obscure the effect size and thus lead to misleading conclusions. Therefore, we decided to investigate the correlation between protein abundance and stability, and its impact on evolutionary sequence constraints using unbinned data from all four species analyzed by Leuenberger *et al.*

We note that despite possible biases and uneven sampling of proteins in different organisms, the correlation of sequence constraints, commonly quantified as the rate of non-synonymous substitutions per site (Ka), with protein abundance (table 1, second column) and gene expression (table 1, third column) remains strong for the subset of proteins with reported T_m_ measurements. Therefore, these data can be used to investigate the nature of sequence constraints in all organisms analyzed by Leuenberger *et al.* Moreover, although proteins with similar T_m_ may have different folding stabilities at physiological temperatures (Becktel and Schellman 1987), using data from the ProTherm database we found a significant correlation between proteins’ T_m_ and their unfolding Gibbs free energies (Spearman’s r=0.64, p<10^-20^, Pearson’s r=0.75, p<10^-20^, supplementary fig. 1). Consequently, reported protein melting temperatures do reflect, at least on average, protein stabilities at physiological temperatures.

Using protein stabilities and abundances from Leuenberger *et al.*, we first confirmed a weak but significant positive correlation between T_m_ and protein abundance in *E. coli* (Spearman’s r=0.16, p=6×10^-6^; Pearson’s r=0.2, p=7×10^-8^). Surprisingly, for the other two organisms with protein abundance data (yeast and human) we found significant negative correlations with T_m_ (Spearman’s r=-0.11 and -0.19, respectively, both p<0.005, Pearson’s r=-0.09 and -0.13, both p<0.02), contrary to the prediction that abundant proteins should be more stable. Moreover, because ribosomal proteins are highly abundant and generally enriched among stable proteins, it is possible that the weak correlation of T_m_ and protein abundance is primarily driven in *E. coli* by the properties of ribosomal proteins. Indeed, excluding 46 ribosomal proteins (out of 730 considered proteins) substantially decreased both the magnitude and the significance of the correlation in *E. coli* (fig 1a; Spearman’s r=0.08, p=0.03, Pearson’s r=0.09, p=0.02), whereas for yeast and human data, we still observed small negative correlations (fig 1b,c, and table 1, fourth column). We next calculated, after removing ribosomal proteins, the correlation between T_m_ and mRNA expression in all four species (fig. 1. d-g, and table 1, fifth column). Similar to protein abundances, and contrary to the expectation of the misfolding avoidance hypothesis, the correlations were either non-significant or negative. Furthermore, when T_m_ was calculated considering data from all peptides associated with each protein, rather than only peptides assigned to the least stable protein domain (the approach used by Leuenberger *et al.* (2017)), we again observed only a weak positive correlation between T_m_ and protein abundance in *E. coli* (Spearman’s r=0.07, p=0.05, Pearson’s r=0.09, p=0.01); but not in any other organism.

**Figure 1.**
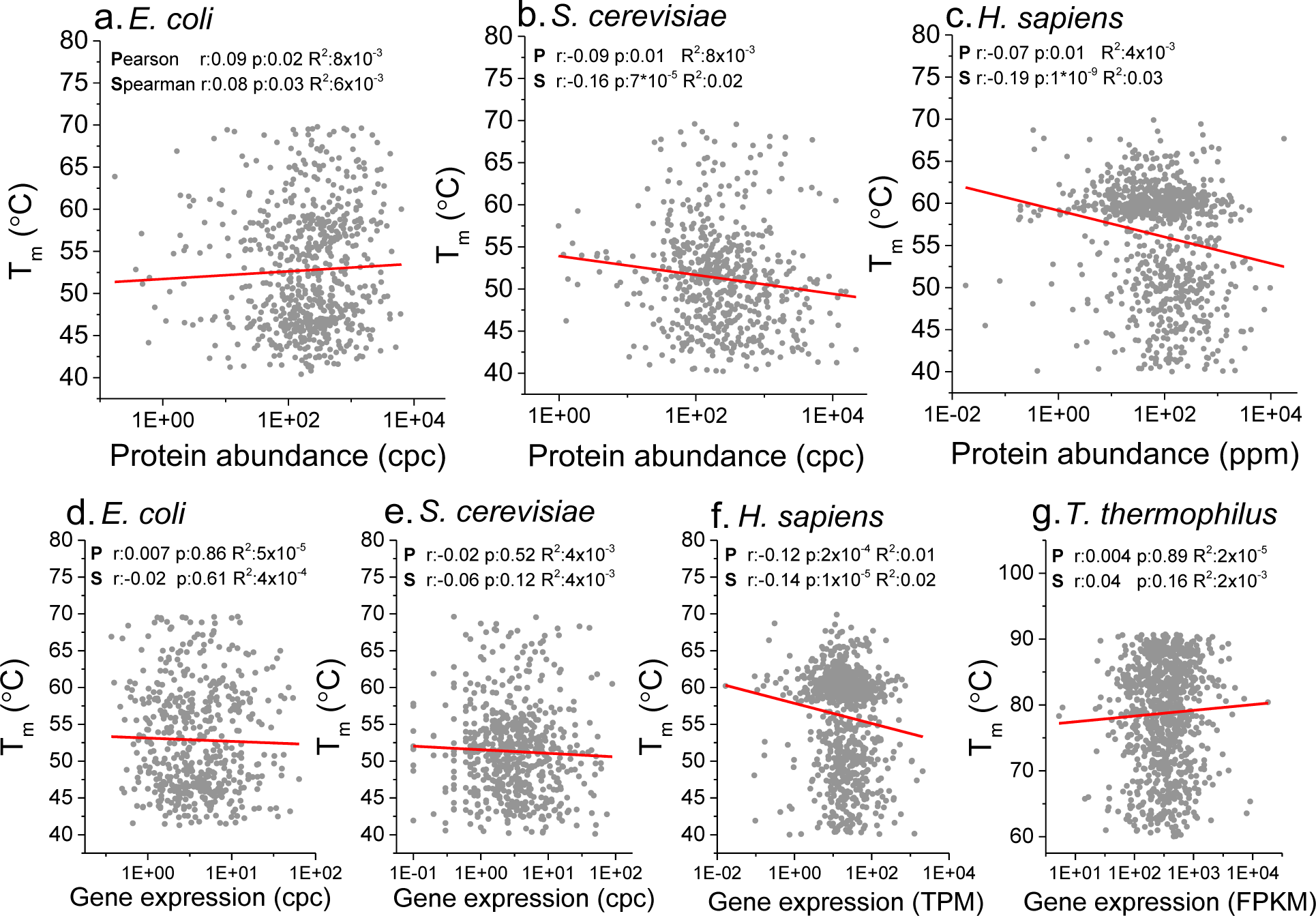
**a-c**. Protein melting temperature (T_m_) calculated by Leuenberger *et al.* as a function of protein abundance in three species (**a.** *E. coli*, **b.** *S. cerevisiae* and **c.** *H. sapiens*). **d-g**. T_m_ as a function of mRNA expression in four species (**d.** *E. coli*, **e.** *S. cerevisiae*, **f.** *H. sapiens* and **g.** *T. thermophilus*). The red lines represent linear fits to the log-transformed protein abundance and mRNA expression data; correlation coefficients, corresponding p-values, and R^2^ are shown for Pearson’s (**P**) and Spearman’s (**S**) correlations in each panel. cpc: counts per cell; ppm: parts per million; TPM: Transcripts Per Kilobase Million; FPKM: Fragments Per Kilobase Million. Proteins annotated as ribosomal were excluded from the analysis.

**Table 1.** Correlation between T_m_, gene and protein expression, and evolutionary rate ^a^

| Species | Protein abundance<br>vs. $K_a^{b,c}$ | Gene expression<br>vs. $K_a$ | $T_m$ vs. protein<br>abundance | $T_m$ vs. Gene<br>expression | $T_m$ vs.<br>$K_a$ |
| --- | --- | --- | --- | --- | --- |
| *E. coli* | -0.38**(-0.38**) | -0.40**(-0.40**) | 0.08* | -0.02 | 0.02 |
| *S. cerevisiae* | -0.47**(-0.47**) | -0.45**(-0.45**) | -0.16** | -0.06 | 0.05 |
| *H. sapiens* | -0.16**(-0.17**) | -0.17**(-0.17**) | -0.19** | -0.14** | 0.02 |
| *T. thermophilus* | N.A. | -0.35**(-0.35**) | N.A. | 0.04 | -0.04 |

<sup>a</sup> Only proteins with measured $T_m$ were considered, ribosomal proteins were excluded (see supplementary table 1 for results including ribosomal proteins); <sup>b</sup> P-values for Spearman's rank correlation are indicated as \* $<0.05$ and \*\* $<5 \times 10^{-3}$ ; <sup>c</sup> Values in parentheses show the partial Spearman correlation between abundance/expression and $K_a$ after controlling for $T_m$

^a^ Only proteins with measured T_m_ were considered, ribosomal proteins were excluded (see supplementary table 1 for results including ribosomal proteins); ^b^ P-values for Spearman’s rank correlation are indicated as * <0.05 and **<5x10^-3^; ^c^ Values in parentheses show the partial Spearman correlation between abundance/expression and Ka after controlling for T_m_

The conjecture that highly expressed proteins are stable because they are designed to tolerate translational errors (Leuenberger et al. 2017) can be directly tested by analyzing the effect of protein stability on the correlation between protein abundance and sequence constraints. Such an analysis demonstrates that the significant negative correlation between protein abundance and evolutionary constraints (Ka), with or without ribosomal proteins, remains essentially unchanged after controlling for protein stability in all analyzed organisms (the correlations in parentheses in the second and third two columns in table 1 and supplementary table 1).

Interestingly, the Leuenberger *et al.* data also suggest that protein stability, irrespective of protein abundance or mRNA expression, does not substantially affect the protein molecular clock. In none of the four species the correlation between T_m_ and Ka is either strong or significant (table 1, last column and fig. 2). There is also no significant correlation between protein stability and the clock rate when only single domain proteins are considered (supplementary fig. 2). These results indicate that, beyond the avoidance of misfolding toxicity, any theory requiring the optimization of protein stability as a dominant constraint of the protein molecular clock is not consistent with the empirical data.

**Figure 2.**
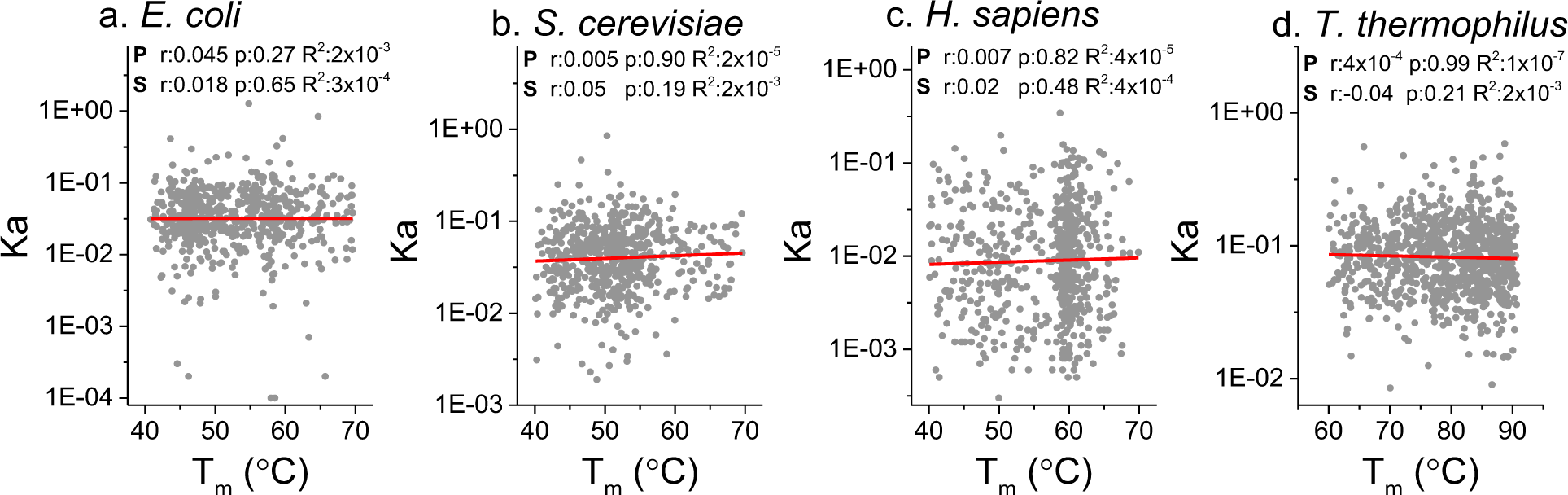
The rate of non-synonymous substitutions per non-synonymous site (Ka) as a function of protein melting temperature (T_m_) calculated by Leuenberger *et al.* (2017). Results are shown for four different organisms (**a.** *E. coli*, **b.** *S. cerevisiae*, **c.** *H. sapiens* and **d.** *T. thermophilus*). Ribosomal proteins were excluded from the analysis. The red lines represent linear fits of the log-transformed Ka data; correlation coefficients, corresponding p-values, and R^2^ are shown for Pearson (**P**) and Spearman (**S**) correlations, in each panel.

Overall, our analyses demonstrate that there is no substantial correlation between protein stability and protein abundance (at most 1-4% of the variance explained). In two of the analyzed organisms, the correlation between stability and abundance is actually opposite to the main prediction of the misfolding avoidance hypothesis. The weak correlation observed in *E. coli* is primarily driven by the properties of ribosomal proteins. There are also no detectable effects of protein stability on the relationships between protein abundance and evolutionary sequence constraints. Therefore, the analysis of the extensive dataset recently generated by Leuenberger *et al.*, similar to previous studies (Plata et al. 2010; Kafri et al. 2016), suggests that neither mistranslation-induced nor spontaneous misfolding toxicity is likely to substantially affect protein sequence constraints and the rate of the protein molecular clock.

Given no significant correlation between T_m_ and Ka, it is likely that common biophysical mechanisms for protein stabilization, such as the burial of several additional hydrophobic residues (Dill and Bromberg 2011), may not significantly increase the evolutionary constraints on hundreds of other sites in a protein. Therefore, it will be important to further investigate in the future how effects associated with the costs of protein production, protein cellular abundance and functional optimization, contribute to evolutionary sequence constraints and the protein molecular clock (Cherry 2010; Plata et al. 2010).

## Materials and methods

T_m_ data, and protein abundances for *E. coli* and yeast, as well as the number of domains per protein, were obtained from supplementary table 3 in the Leuenberger *et al.* study (2017). Human protein abundances were obtained from the whole organism integrated dataset in the PaxDB v.4 database (Wang et al. 2012). *E. coli, T. thermophilus* and *S. cerevisiae* expression data were obtained from Lu *et al.* (2007), Swarts *et al.* (2015) and Holstege *et al.* (1998), respectively. Human expression data were averaged across the main 9 tissues in the Melé *et al.* (2015)’s study. Ka values for *E. coli, S. cerevisiae, H. sapiens* and *T. thermophilus* were calculated with the PAML package (Yang 1997) relative to *Salmonella enterica, Saccharomyces bayanus, Macaca mulatta*, and *Thermophilus aquaticus* orthologs, respectively. Orthologs were identified as bi-directional best hits (BBHs) using protein BLAST (Altschul et al. 1997); we only considered for the analysis BBHs for which at least 70% of the length of the shortest protein was aligned. Unfolding free energy and melting temperature data used in supplementary figure 1 were obtained from the ProTherm (Feb. 2013) database (Bava et al. 2004). The Ribosomal Protein Gene Database was used to identify ribosomal proteins (Nakao et al. 2004).

## Supplementary Tables and Figures

**Supplementary table 1.** Correlation between T_m_, gene and protein expression, and evolutionary rate ^a^ including ribosomal proteins.

| Species | Protein abundance<br>vs. $K_a^{b,c}$ | Gene expression<br>vs. $K_a$ | $T_m$ vs. protein<br>abundance | $T_m$ vs. Gene<br>expression | $T_m$ vs.<br>$K_a$ |
| --- | --- | --- | --- | --- | --- |
| *E. coli* | -0.47**(-0.46**) | -0.50**(-0.49**) | 0.16** | 0.11* | -0.08* |
| *S. cerevisiae* | -0.51**(-0.51**) | -0.53**(-0.53**) | -0.11* | -0.01 | 0.02 |
| *H. sapiens* | -0.18**(-0.18**) | -0.19**(-0.19**) | -0.19** | -0.13** | 0.02 |
| *T. thermophilus* | N.A. | -0.39**(-0.39**) | N.A. | 0.09** | -0.08* |

<sup>a</sup> Only proteins with measured $T_m$ were considered; <sup>b</sup> P-values for Spearman's rank correlation are indicated as \* <0.05 and \*\* <5×10<sup>-3</sup>; <sup>c</sup> Values in parentheses show the partial Spearman correlation between abundance/expression and $K_a$ after controlling for $T_m$

^a^ Only proteins with measured T_m_ were considered; ^b^ P-values for Spearman’s rank correlation are indicated as * <0.05 and **<5x10^-3^; ^c^ Values in parentheses show the partial Spearman correlation between abundance/expression and Ka after controlling for T_m_

**Supplementary figure 1.**
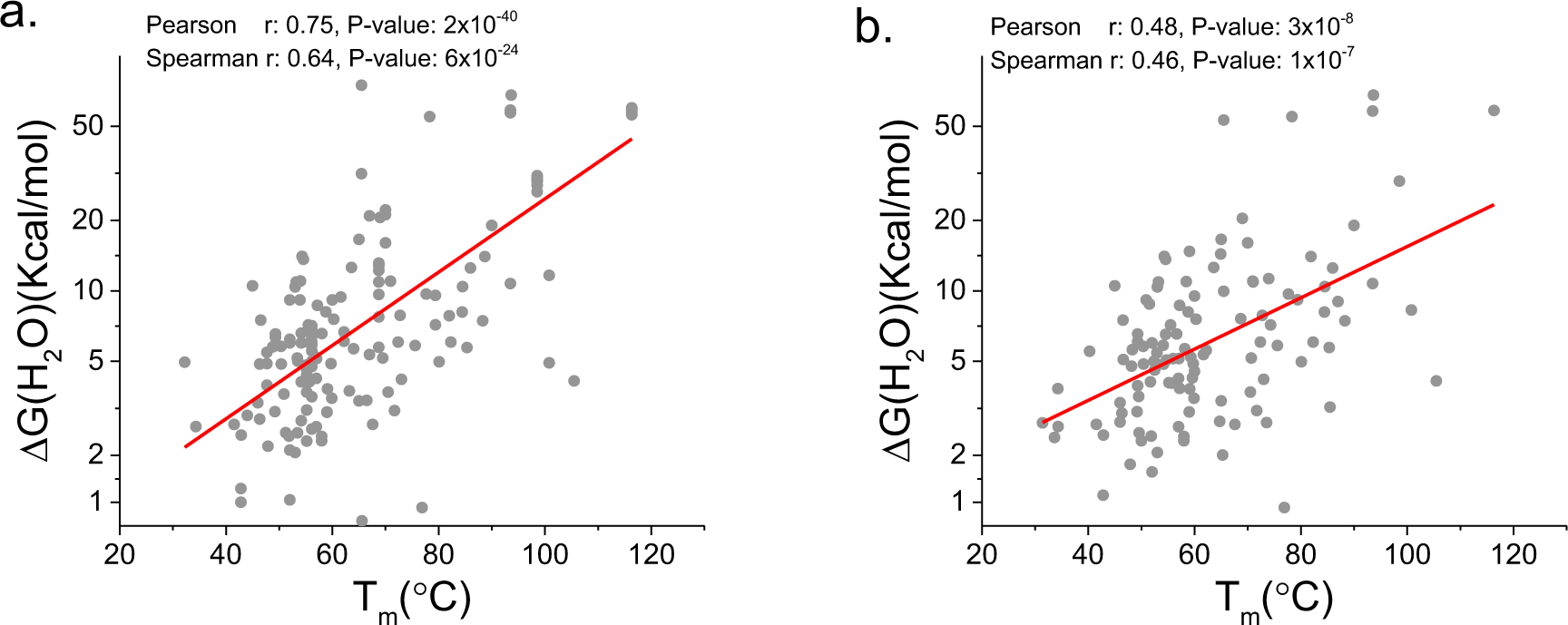
The correlation between melting temperature (T_m_) and folding free energy change (ΔG(H_2_O)) for wild-type proteins in the ProTherm database (Bava et al. 2004). **a.** The correlation between T_m_ and ΔG(H_2_O) across proteins and experimental conditions (n=164); each point corresponds to values of ΔG(H_2_O) and T_m_ of a protein measured at a specific temperature and the same pH. **b.** The correlation between T_m_ and ΔG(H_2_O) across different proteins (n=117). Each point represents one protein, with values of Tm and ΔG(H_2_O) averaged across experimental pH and temperature conditions. The red lines represent a linear fit to the log-transformed ΔG(H_2_O) data.

**Supplementary figure 2.**
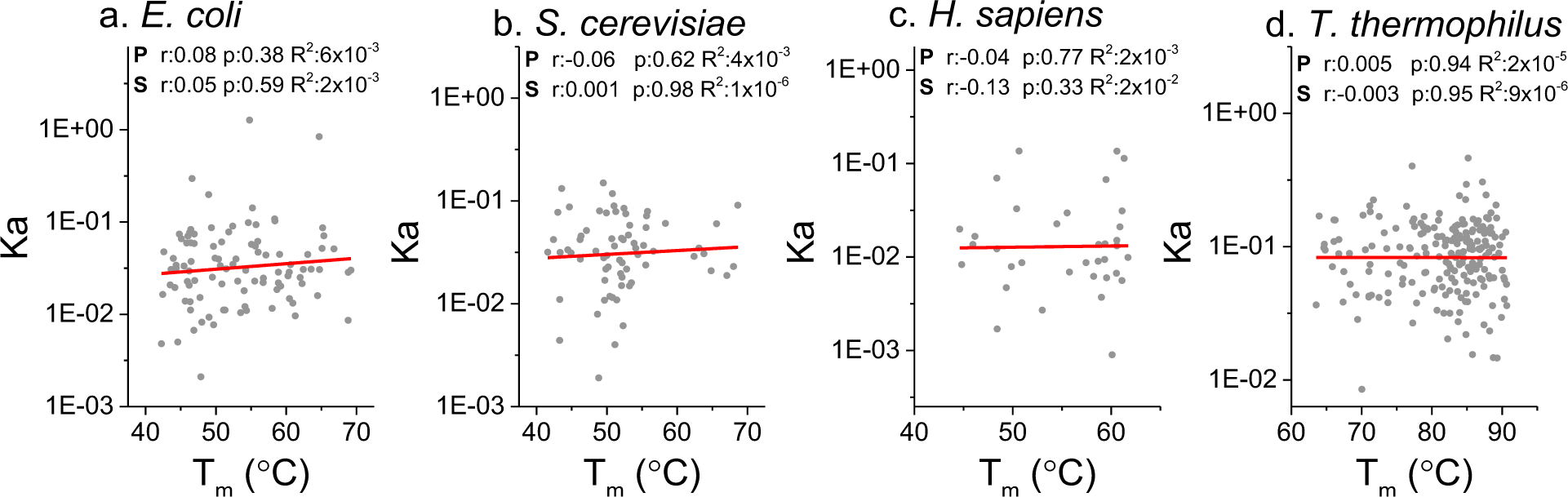
The rate of non-synonymous substitutions per non-synonymous site (Ka) as a function of protein melting temperature (T_m_) for single domain proteins. Results are shown for four different organisms (**a-d**). Single domain proteins were defined as those with a single predicted pfam domain and a single measured domain according to Leuenberger et al. (2017); based on the hierarchical clustering of T_m_ values across measured peptides. Ribosomal proteins are excluded from the analysis. The red lines represent linear fits of the log-transformed Ka data; correlation coefficients, p-values, and corresponding R^2^ are shown for Pearson (P) and Spearman (S) correlations, in each panel.

